# Protective effect of prior knowledge in ageing

**DOI:** 10.64898/2025.12.04.692350

**Authors:** Georgia Milne, Oris Shenyan, Laura Haye, Matteo Lisi, Tessa Dekker

## Abstract

Visual perception depends on both sensory input and prior knowledge, and the quality of these information sources changes across the lifespan. As people age, accumulated knowledge, and its influence on perception, tend to increase, while declines in low-level visual functions reduce input quality and bias perception toward prior information. Yet, the combined effects of these changes on higher-level visual perception remain unclear, partly because concurrent cognitive decline may complicate the use of stored knowledge. In this study, we examined how prior knowledge shapes visual perception in younger (<40 years) and older (>60 years) adults using ambiguous two-tone images, whose recognition depends on prior knowledge. Providing participants with the original photographs used to generate the two-tones triggered perceptual reorganisation, improving recognition in both groups. Although older adults recognised fewer two-tones overall - a pattern linked to reduced visual acuity - they showed a comparable benefit from prior knowledge when cued. These findings suggest that, despite age-related sensory decline, the ability to draw on prior knowledge remains robust in older adulthood and may help preserve perceptual function.

## Introduction

Human perception is a dynamic process that integrates sensory input with internally generated predictions. From a Bayesian perspective, the brain balances these two sources of information according to their reliability: when inputs are clear and informative, perception depends heavily on them, whereas in noisy or ambiguous scenes, prior knowledge exerts greater influence (Lee & Mumford, 2003). The information available to the brain, however, changes across the lifespan. Ageing is accompanied by declines in sensory quality, alongside an accumulation of knowledge and experience. At the same time, changes in memory and cognition may limit access to this knowledge. This study therefore examines how age-related changes in sensory input and knowledge access jointly shape how we perceive the world as we age.

In later life, we typically experience changes across visual and cognitive functions (Nagarajan et al., 2022). Age-related declines of basic sensory and perceptual processes are varied and well-documented. With age, our visual acuity, contrast sensitivity, motion discrimination, and other low-level visual functions diminish (Andersen, 2012; Faubert, 2002; Owsley, 2011). Cognitive abilities, including the capacity to encode, retain, and retrieve information, also tend to decline with age (Cadar et al., 2018; Craik, 2023). These changes do not occur in isolation; visual impairment is a strong predictor of this decline (Eppenberger et al., 2024; Reyes-Ortiz et al., 2005). This correlation may reflect shared risk factors, increased demands on memory and attention due to degraded inputs, or reduced engagement in stimulating activities following vision loss (Eppenberger et al., 2024; Nagarajan et al., 2022; Whitson et al., 2018; Zheng et al., 2018).

Meanwhile, object knowledge is well-established in old age, and there is growing evidence that older adults may rely more heavily on their prior knowledge, which can enhance their performance in certain perceptual contexts. Behavioural studies have revealed age-related biases toward prior knowledge in visual search (Wynn et al., 2020), illusory perception (Botwinick et al., 1959; Dowlati et al., 2016; Mazuz et al., 2024), speech perception (Sheldon et al., 2008), and memory tasks (Badham et al., 2016; Loaiza et al., 2015), for example. Complementing these findings, neuroimaging studies suggest increased reliance on feedback and top-down processes. Older adults show reduced activation in sensory regions associated with bottom-up input, alongside greater activation in frontoparietal regions linked to top-down control (Davis et al., 2008; Dowlati et al., 2016; Lai et al., 2020).

Taken together, these findings raise the possibility that reduced reliability of sensory inputs, despite accompanying changes in some cognitive domains, may be offset by a preserved or even enhanced ability to rely on prior knowledge. To examine this, and to test whether such reliance helps preserve perceptual function in later life, we measured older adults’ perception of two-tone images - stimuli whose recognition critically depends on prior knowledge (Figure 1A).

**Figure 1:**
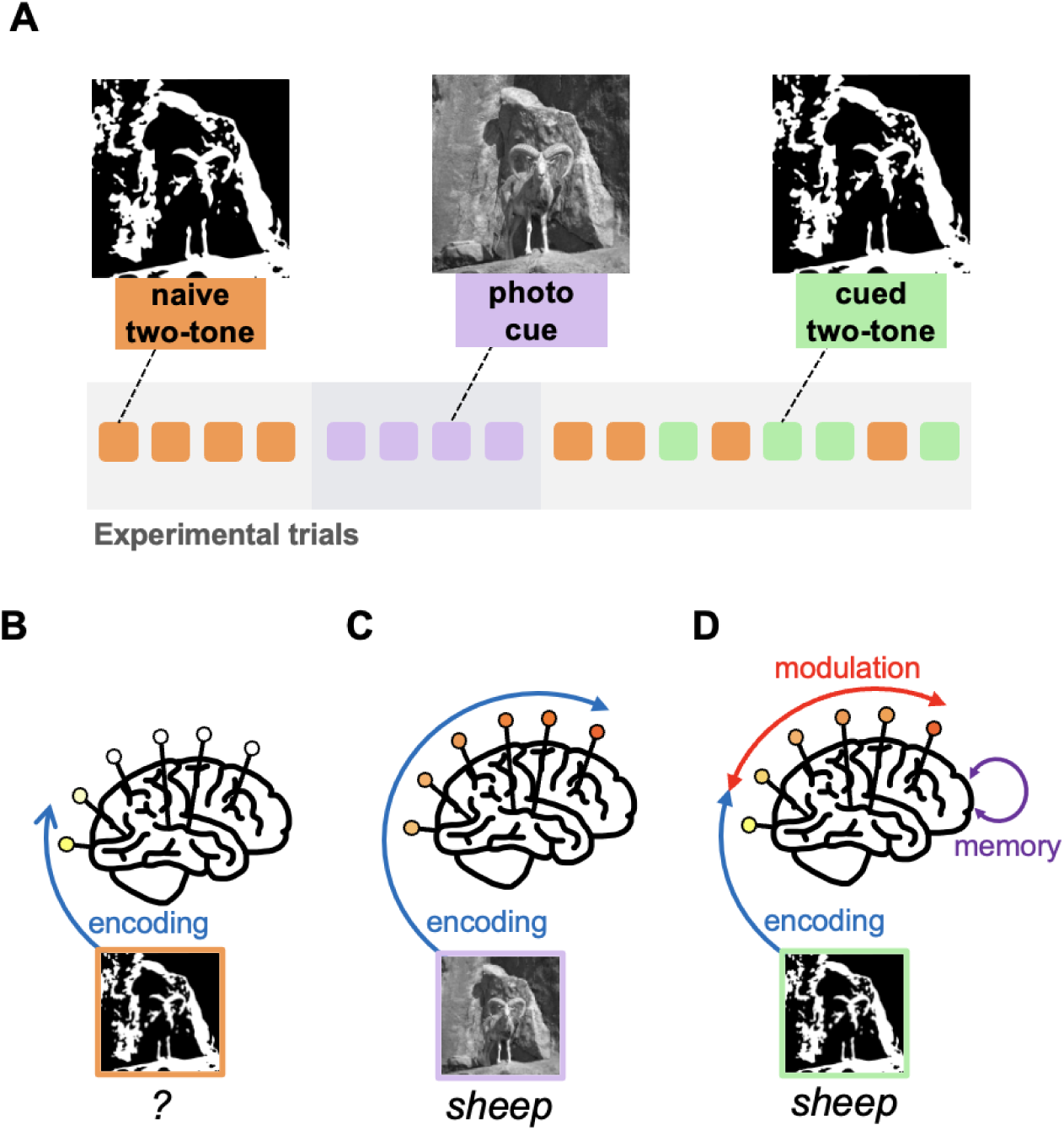
Experimental procedure and mechanisms of two-tone perception. **A** Experimental procedure: Each image was presented for 200ms in three conditions, as a naive two-tone (orange), then as a photo cue (lilac), and finally as a cued two-tone (green). Naive and cued two-tone conditions of different images were shuffled together, and always presented before (naive) and after (cued) the corresponding photo condition. **B-D** Mechanisms of two-tone perception: two-tone images do not include sufficient visual information to be recognised naively (**B**), photo cues are recognised immediately via bottom-up processing, activating higher-order concept areas (**C**), after cueing, the same two-tone can be easily recognised due to the integration of bottom-up processing and top-down signals containing prior knowledge (**D**).

First popularised in the field of psychology by Craig Mooney (Mooney, 1957), two-tone images are black-and-white photos that have been distorted through Gaussian smoothing and binarisation. These distorted versions are difficult to recognise when viewed naively (Figure 1B). However, after seeing the undistorted photo version of these images (Figure 1C), subsequent recognition of the two-tone version becomes automatic (Figure 1D). This process, known as perceptual reorganisation, is attributed to the incorporation of prior knowledge with incoming sensory information via top-down pathways (Friston & Kiebel, 2009; Rauss & Pourtois, 2013). Evidence suggests that feedback from higher-order cortical regions, such as the lateral occipital complex and prefrontal cortex, to lower-level visual areas plays a crucial role in the recognition of two-tone images in young adults (Bona et al., 2016; van Loon et al., 2016). Two-tone cueing paradigms, where prior knowledge is introduced via informative visual cues, therefore provide a useful paradigm for studying the influence of top-down processing. In previous work, we demonstrated that young children show a drastically reduced knowledge-guided benefit when parsing cued two-tones at the level observed in adults, with a large developmental shift in the ability to recognise these stimuli during late childhood (Milne et al., 2024). These findings show that the ability to use prior knowledge to recognise ambiguous visual objects offers a challenge to the early developing system. It is unclear how this ability is impacted in older age, when knowledge and neural connections across visual brain regions are long-established, but visual and cognitive skills may decline.

Altered perception of two-tone images has been found during adulthood in pathological contexts. For example, individuals with psychosis, as well as psychosis-prone healthy individuals, display a greater perceptual reorganisation effect than control groups (Teufel et al., 2015). The authors of this study suggest that participants may upweight prior knowledge to account for increased sensory noise, perhaps due to psychosis-related deficits in early visual processing, or the various visual deficits associated with schizotypy disorders (Adámek et al., 2022; Kéri et al., 2002; Shoham et al., 2023). Patients with Lewy body disease who experience visual hallucinations have also been found to experience increased perceptual reorganisation of two-tone images compared to those without hallucinations (Zarkali et al., 2019). This was attributed to impairments in early sensory areas leading to a loss of feedforward signal (i.e., less reliable inputs) and a compensatory upweighting of feedback signals conveying prior knowledge. In both these cases, increased reliance on prior knowledge in perception was attributed to reduced sensory input quality. One possibility therefore is that the reductions in sensory signal reliability that occur during healthy ageing (affecting encoding in Figure 1B), combined with the well-established knowledge and neural network connections at this stage of life, may result in an upweighting of priors in healthy older adults (enhanced knowledge-driven recognition, Figure 1D).

Alternatively, older adults may exhibit decreased patterns of perceptual reorganisation compared to younger adults, similar to the pattern we previously observed in children (Milne et al., 2024). Age-related declines in visual function may affect the ability to encode visual information, impacting not only the perception of the two-tone stimuli but also that of the photo cue (Figure 1C). This may therefore hinder the robust activation of knowledge representations needed for reorganisation. In addition, impairments in memory that occur in later years of life (Cadar et al., 2018; Craik, 2023) may limit the ability to access and maintain the prior knowledge of image content necessary to parse the cued two-tone (Figure 1D). Therefore, the net effect of sensory and cognitive declines as we age could also manifest as decreased levels of perceptual reorganisation.

Taken together, these findings raise the possibility that reduced reliability of sensory inputs, despite accompanying changes in some cognitive domains such as memory, may be offset by a preserved or even enhanced ability to rely on prior knowledge. To examine this possibility, and to test whether such reliance helps preserve perceptual function in later life, we measured older adults’ perception during a two-tone cueing task, an established measure of the perceptual influence of prior knowledge.

## Methods

### Participants

Forty-seven participants completed the study. Two participants were excluded due to poor visual acuity (≥ 0.4 logMAR). Of the remaining 45 participants, 24 were in the younger age group (15 female, 9 male, mean age: 27.25 years, SD: 5.45, age range: 19-39), and 21 were in the older age group (11 female, 10 male, mean age: 68.9 years, SD: 5.63, age range: 60-79). Participants had no reported history of eye disease, or developmental or neurological disorders. Participants were recruited through the UCL Psychology Subject Pool (‘SONA’), the NIHR Join Dementia Research database and word of mouth. Participants received £10/hour compensation. All participants provided informed consent. The experimental procedure was approved by the Research Ethics Committee at University College London. The research was carried out in accordance with the tenets of the Declaration of Helsinki.

### Experimental paradigms

#### Visual and Cognitive function

##### Contrast sensitivity

Contrast sensitivity (CS) was assessed via a binocular Mars letter contrast sensitivity test (Mars Perceptrix, Chappaqua, NY; http://www.marsperceptrix.com/). This is a portable chart with eight rows of six letters of constant size in each row (48 letters in total), which decrease in contrast at a rate of 0.04 log unit steps. The test was administered at 50 cm and illuminated by a lamp. Tests were scored with a value of 0.04 log CS per letter named correctly, with a maximum possible score of 1.92 log CS.

##### Visual acuity

Visual acuity (VA) was assessed via a Good-Lite mini Snellen chart (Good-Lite, Elgin, IL; https://good-lite.com/), a compact, portable version of a traditional Snellen chart. The test was administered at 40 cm and illuminated by a lamp. Eye correction was measured in LogMAR units, ranging from 1.4 (worst) to -0.1 (best). Participants were asked to wear any eye correction utilised on a day-to-day basis while completing the test.

##### Short-term image memory

To measure task-relevant image memory, we took data from an inhibitory control task (The Inhibition of Current Irrelevant Memories task [ICIM; Alderson-Day et al., 2019]). The task consists of three blocks of 95 sequentially presented black-and-white line drawings (Snodgrass & Vanderwart, 1980). Each block includes the same 60 unique images (40 pictures presented once, 5 pictures presented twice and 15 pictures presented three times. Stimuli are presented for 2000 ms, followed by a 700 ms interstimulus interval during which participants respond to whether each image was a first presentation or a repeat within the current block via a button press ( “1” for first, “2” for repeat). In the second and third blocks, participants distinguish whether a repeat was completely novel (press “1”), presented in a previous block (press “1”, inhibiting an irrelevant memory of seeing the object before), or presented before in this block (press “2”). To measure image memory as relevant to our two-tone test of perceptual reorganisation, we measured the number of repeated images that participants failed to recognise as a repeat (‘misses’). This offers an index of short-term image memory across an approximately matched timespan as the repeated image presentations in our two-tone task.

#### Two-tone task

##### Apparatus

Stimuli were presented on a 32″ LED backlit LCD monitor (60 Hz frame rate; Cambridge Research Systems BOLDscreen) driven by an Apple MacBook Pro, running MATLAB R2015b with the Psychophysics Toolbox (2015).

##### Stimuli

Stimuli were generated from 24 images of animals and objects from The Berkeley Segmentation Dataset and Benchmark (Martin et al., 2001) using Matlab R2022a (2022). Images were converted to greyscale and cropped to a square to create photo cues. Two-tone stimuli were created using methods previously described elsewhere (Milne et al., 2024). Specifically, photo cues were smoothed with a 2D Gaussian filter and then thresholded to binarise pixel luminance to black or white, creating images with obscured edges. Smoothing levels and binarisation thresholds varied per image; standard deviation of Gaussian distribution ranged from 0.5 - 6.0 SD, binarisation thresholds ranged from 0.4 - 2.0 times the mean luminance of the greyscale image. Six of these two-tones were generated with minimal smoothing, of which four were used in practice trials and two were used as catch trials in the main task to promote motivation and to obtain an index of attentiveness and task comprehension. All stimuli were presented as a 13 degrees of visual angle square centred on a 50% grey background.

##### Procedure

Participants sat ∼70 cm in front of a 32″ LED backlit LCD monitor (60 Hz frame rate; Cambridge Research Systems BOLDscreen) in a darkened and soundproofed room. The experimenter was present throughout the procedure. Following task instructions, participants completed a training task in which a greyscale image was displayed and transformed gradually into a two-tone image that was unsmoothed and easy to recognise. To confirm that participants understood that image content was maintained across the greyscale and two-tone, they were asked to indicate whether the two-tone was an object or an animal. Following this, participants completed four practice trials with unsmoothed two-tones. This was followed by the main task, in which 20 two-tones of varying difficulty and the corresponding 20 greyscale images were presented in a pseudo-randomised order, such that for each greyscale image the corresponding two-tone was presented twice - once preceding the greyscale image (naive condition), and once following the greyscale image (cued condition). The task included 60 experimental trials in total, with each trial consisting of a fixation cross presented for 1 s, followed by either a naive, greyscale or cued image presented for 200 ms, and finally a response screen. During the response screen, the question ‘Can you name what you saw?’ was displayed, prompting the participant to verbally name what they perceived in the image. The given answer was then typed in by the experimenter and displayed on the screen. Motivational but uninformative feedback was given after each trial to all participants. A break screen was displayed every 12 trials to show progress through the task. Including these five short breaks, the task lasted approximately 15 minutes in total.

##### Scoring

To quantify image recognition, we measured naive, greyscale and cued naming accuracy. Image names were scored as correct if the content was correctly identified at the basic category level - superordinate category names (e.g., naming a cow an ‘animal’) were scored as incorrect; however, simple basic level or subordinate categories were accepted (e.g., a sheep named as ‘goat’, or scissors named as ‘shears’) as long as there was consistency in naming across the two-tone and greyscale conditions for each participant. Two experimenters independently scored each image and met to resolve any discrepancies between scorings. For all further analyses, trials of any images that were answered incorrectly in the greyscale condition were removed for each participant. Our primary outcome measure, the cueing effect, was quantified as the difference between the number of cued correct images and naive correct images, excluding trials where the grayscale image was named incorrectly.

### Statistical analysis

We adopted two complementary analytical approaches to assess group and condition differences in task performance.

First, we used generalised linear mixed-effects models (GLMMs), which provide a flexible framework for analysing binary outcome data while accounting for the non-independence of observations. Recognition accuracy (correct = 1, incorrect = 0) was modelled using a binomial distribution with a logit link function, as implemented in the lme4 package in R (Bates et al., 2015). This approach allowed us to account for unequal trial numbers across participants (due to the removal of images for which the photo cue was unrecognised), as well as random variation associated with individual participants and image items.

We fitted separate models for the three different image conditions (photo cue, naive two-tone and cued two-tone), each including age group as a fixed-effect predictor of recognition accuracy, with random intercepts for participant and image. To assess the effects of visual and cognitive abilities on task performance, we fitted further GLMMs including either visual acuity, contrast sensitivity, or short-term image memory as a fixed-effect predictor of performance for each of the three image conditions, with the same random intercepts as previously.

To quantify the cueing effect, we fitted a further GLMM including fixed effects of age group, two-tone condition (naïve vs. cued), and their interaction, with the same random effects structure. Finally, to test whether visual function moderated the cueing effect, we added visual acuity and its three-way interaction with condition and age group as fixed effects. Model coefficients were converted to odds ratios (ORs) for interpretability, with values below 1 indicating lower odds of correct recognition in older versus younger adults.

As a complementary, assumption-light approach, we conducted Mann–Whitney U tests to compare performance between younger and older adults on overall visual and cognitive task measures, as well as within each image condition (naïve, photo cue, cued) and for the cueing effect. This non-parametric test offers an intuitive comparison of central tendencies and serves as a robustness check on the model-based analyses.

All correlations reported are Spearman’s rank correlations, given the non-normal distribution of several variables. Statistical significance was defined as p < .05. Full model outputs and descriptive statistics are reported in the Results section.

## Results

We tested the ability to use prior knowledge to perceive hard-to-recognise two-tone images in 24 young adults (19 - 39 years) and 21 older adults (60 - 79 years). Participants completed a two-tone cueing task, where they were asked to name the content of photos each presented for 200 ms, first as two-tone images (naive two-tones), then as the original greyscale photos (cues) and finally as two-tones again (cued two-tones). In addition, participants were also tested on visual and cognitive functions that may affect their performance on this task, including visual acuity, contrast sensitivity and short-term image memory.

### Photo recognition

We first considered the ability of younger and older participants to recognise images presented as undistorted greyscale photos (cues). Across the two age groups, cue recognition accuracy was high (>90%) and did not significantly differ (Mann-Whitney U test: *p* > 0.05). This was further confirmed by a generalised linear mixed effects model with age group as a fixed-effect predictor for photo recognition, which showed no significant effect of age group on cue recognition scores (OR = 0.31; SE = 0.21; Z = -1.69; *p* = 0.092). This suggests good understanding and engagement with the task across both groups, and no large age differences in participants’ ability to recognise and name the content of the images in the task.

### Two-tone recognition

Next, we assessed participants’ ability to recognise images presented as two-tones both before and after viewing an informative photo cue, and calculated the perceptual benefit afforded by photo cueing. Compared to younger adults, older adults were significantly less able to recognise the content of naive two-tones (two-tones viewed before photo cueing; Figure 2A; Mann-Whitney U test: *p* = 0.028, PS = 0.31). This age effect was confirmed with a generalised linear mixed effects model, which showed a significant effect of age group on naive two-tone recognition scores (OR = 0.43; SE = 0.16; z = -2.33; *p* = 0.020).

**Figure 2:**
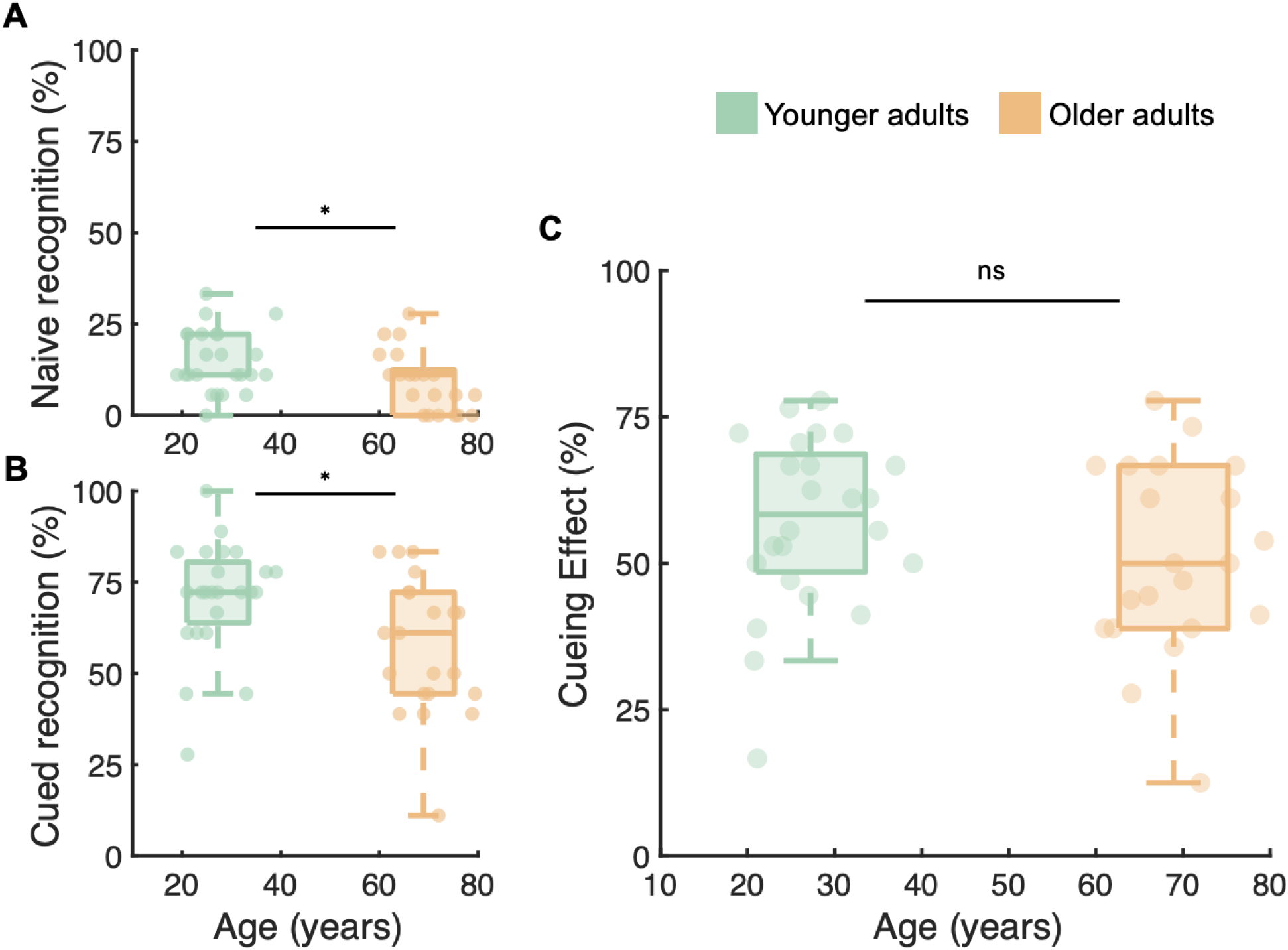
Two-tone recognition across age. **A** Naive two-tone recognition for younger (green) and older (orange) populations. * denotes Mann-Whitney U significance of *p* < 0.05. **B** Cued two-tone recognition for younger (green) and older (orange) populations. * denotes Mann-Whitney U significance of *p* < 0.05. **C** Cueing effect ([cued recognition - naive recognition] / photo recognition) for younger (green) and older (orange) populations. ns denotes non-significance (Mann-Whitney U significance of *p* > 0.05).

After viewing the undistorted photo cues, recognition of the same two-tone images significantly increased for both age groups (Mann-Whitney U test, younger adults: *p* < 10^-5^, PS = 0.003; older adults: *p* < 10^-5^, PS = 0.018), revealing large cueing effects for both younger and older adults. Despite this, the benefit of cueing was not sufficient to overcome the initial differences found in two-tone recognition across age groups. As with naive two-tones, younger adults were able to recognise more cued two-tones than older adults (Figure 2B; Mann-Whitney U test: *p* = 0.014, PS = 0.29). This finding was further supported by a generalised linear mixed effects model, which showed a significant effect of age group on cued two-tone recognition scores: OR = 0.45; SE = 0.15; Z = -2.40; *p* = 0.016).

To compare the size of the cueing effect on two-tone recognition across age, we calculated the recognition improvement between naive and cued two-tone conditions for all images that were correctly recognised in the photo condition. Despite poorer performance for both naive and cued conditions, older adults received an equal benefit from viewing the photo cue as did younger adults (Figure 2C; Mann-Whitney U test: *p* > 0.05). We also ran a general linear mixed effects model with two-tone condition (cued or naive) and the interaction between age group and condition as fixed-effects predictors of recognition of two-tone images. This revealed main effects of age group and condition on overall accuracy, but no significant interaction effect between age group and condition (OR = 0.99; SE = 0.32; Z = -0.04, *p* = 0.97), again suggesting that older adults experience a similar cueing effect to younger adults. An equivalence test further indicated that we can reject the presence of interaction effects beyond a small-to-medium effect size (approximate Cohen’s *d* = 0.35; p = 0.029).

### Visual and cognitive function

In addition to the two-tone cueing task, participants also completed tests of visual and cognitive functions that may affect their ability to encode, retrieve, and maintain the content of two-tone and photo stimuli.

We tested participants’ short-term visual memory using the Inhibition of Current Irrelevant Memories task (ICIM; Alderson-Day et al., 2019). This task was one of a cohort of tests completed as part of a separate analysis, however the overall miss rate (repeated images that were not identified as having been seen before) provides a useful measure of image memory over a comparable length of time as in the two-tone cueing task. Here, we found no significant differences between the miss rates of younger and older adults (Figure 3A; Mann-Whitney U test: *p* > 0.05), suggesting that reduced photo-cue recall is unlikely to have limited the cueing effect in the older group.

**Figure 3:**
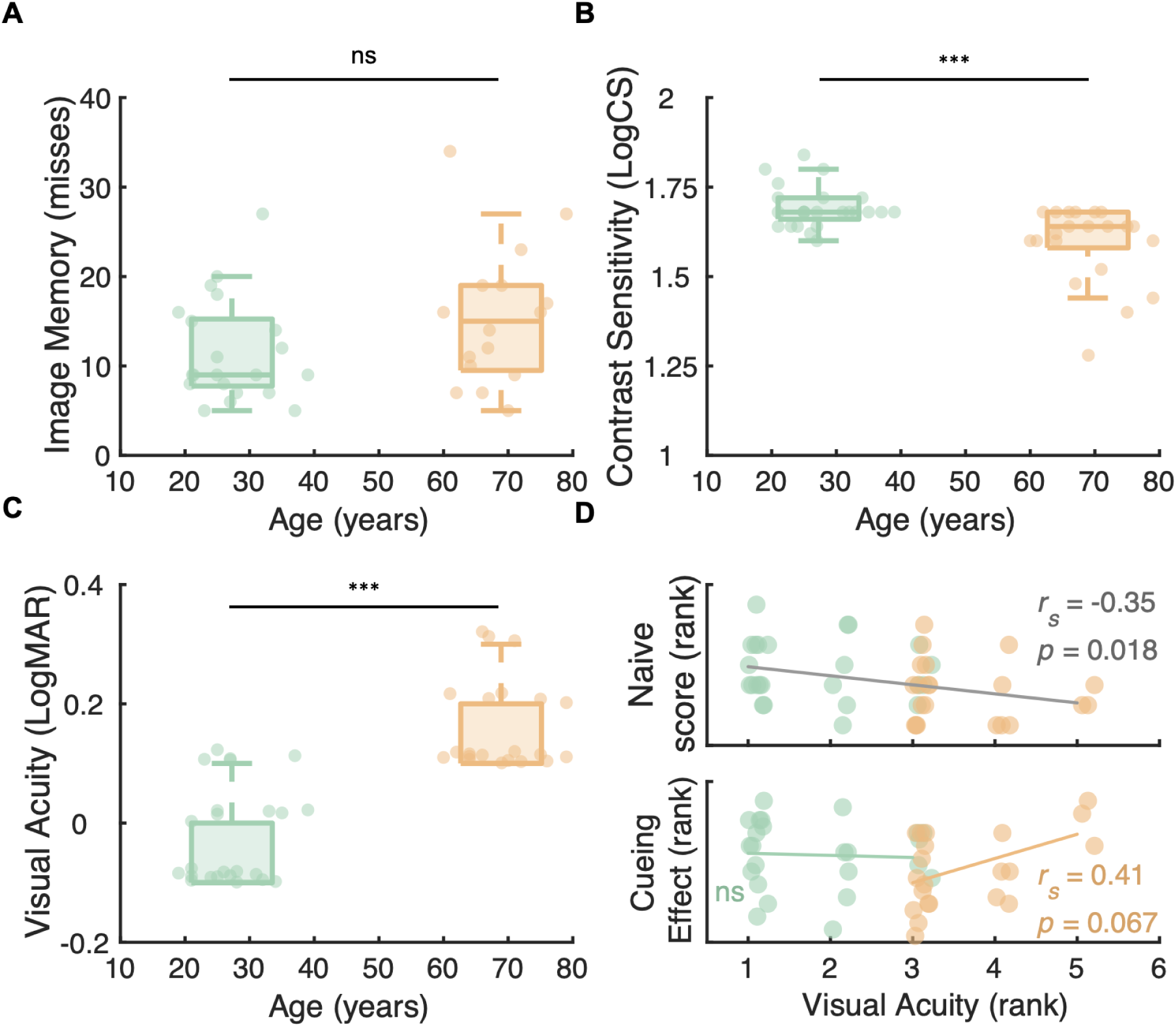
Visual and cognitive function measures across age. **A** Short-term image memory performance for younger (green) and older (orange) populations, ns denotes Mann-Whitney U significance of *p* > 0.05. **B** Contrast sensitivity of younger (green) and older (orange) populations, *** denotes Mann-Whitney U significance of *p* < 10^-3^**. C** Visual acuity of younger (green) and older (orange) populations, where higher LogMAR values indicate poorer visual acuity. *** denotes Mann-Whitney U significance of *p* < 10^-3^**. D** Top panel: Rank naive two-tone performance and rank visual acuity scores, grey line shows least-squares linear trendline, and r and *p* values correspond to a Spearman’s rank correlation across both age groups. Bottom panel: Rank cueing effect and rank visual acuity scores, coloured lines show least-squares linear trendline for younger (green) and older (orange) participants. r and *p* values correspond to a Spearman’s rank correlation within older participants, ns denotes Spearman’s rank significance of *p* > 0.05.

We next tested participants’ contrast sensitivity and visual acuity, and found large differences in these visual abilities across the two age groups. Older adults had significantly poorer contrast sensitivity (Figure 3B; Mann-Whitney U test: *p* < 10^-3^, PS = 0.18), and while the average best-corrected visual acuity measure for younger adults equated to better than 20/20 vision, older adults’ visual acuity was significantly poorer (Figure 3C; Mann-Whitney U test: *p* < 10^-5^, PS = 0.064).

Differences in these visual abilities were found to impact performance on the two-tone cueing task in distinct ways. Across both age groups, participants with better contrast sensitivity achieved higher recognition scores for photo cues (Spearman’s rank: ***r****_s_*(43) = 0.43, *p* = 0.003). This link was further substantiated by a general linear mixed effects model with contrast sensitivity as a fixed-effect predictor of cue recognition (OR = 2.23; SE = 0.67; Z = 2.68, *p* = 0.007). However, we found no significant effects of contrast sensitivity on either naive or cued two-tone recognition scores, or cueing effect (the improvement in recognition between these conditions).

Participants across both age groups with better visual acuity scores achieved higher recognition scores for naive two-tone images (Figure 3D, top panel; Spearman’s rank: ***r****_s_*(43) = -0.35, *p* = 0.018). This effect was confirmed by a generalised linear mixed effects model including visual acuity as a fixed-effect predictor of naive recognition score (OR = 0.65; SE = 0.12; z = -2.42; *p* = 0.016). No equivalent effect of visual acuity was found for recognition of the same two-tone images viewed after photo cueing (Spearman’s rank: *p* > 0.05; generalised linear mixed effects model including visual acuity as a fixed-effect predictor of cued recognition score [OR = 0.12; SE = 0.17; z = -1.47; *p* = 0.14]), suggesting that prior knowledge granted by the photo cue is sufficient to overcome this acuity-driven deficit in two-tone recognition.

In fact, for older adults, we found a non-significant trend of larger cueing effects experienced by those with worse visual acuity scores (Figure 3D, lower panel; Spearman’s rank: ***r****_s_*(19) = 0.41, *p* = 0.067). This trend was not present amongst the younger adults (Spearman’s rank: ***r****_s_*(22) = -0.12, *p* = 0.57). Evidence for this age-dependent pattern was provided by a general linear mixed effects model including visual acuity, age group, two-tone condition (cued or naive), and the interaction between visual acuity, age group, and condition as fixed-effect predictors for overall two-tone accuracy. Here, a significant three-way interaction effect between acuity, age group and condition (OR = 2.93; SE = 1.57; Z = 2.02, *p* = 0.044) suggests that visual acuity may differentially affect the perceptual benefit received from cueing across age groups. Specifically, it may indicate that people with lower acuity in old age show stronger cueing effects, but that this relationship is not present in younger adulthood when acuity is generally better and more consistent. As our sample sizes are small and correlations between acuity and perceptual reorganisation are only trending within the older age group (p = 0.067), we consider this a preliminary finding. Nevertheless, this result aligns with the proposed Bayesian reconciliation of visual input and prior knowledge for two-tone recognition, where diminishing quality of bottom-up input sources during old age may result in an upweighting of priors.

## Discussion

Using a two-tone cueing paradigm, we examined the influence of prior information on the perception of impoverished visual stimuli, and how this varied with ageing. To trigger perceptual reorganisation - the automatic recognition of a stimulus after acquiring relevant information - we presented participants with difficult-to-recognise, binarised images (naive two-tones) before presenting the corresponding undistorted photo version (photo cue) and finally the same two-tone version again (cued two-tones). We show that while older adults exhibited reduced performance for both naive and cued two-tone recognition, they showed no differences in perceptual reorganisation compared to younger adults. This suggests that despite large reductions in visual function compared to the younger cohort, older adults were equally able to use photo cueing to facilitate the perception of degraded visual scenes.

As anticipated, older participants exhibited significantly poorer basic vision levels than younger adults. Our sample of older adults (>60 years) scored an average of 0.15 logMAR visual acuity and 1.59 log contrast sensitivity, whereas our younger cohort achieved better than 20/20 vision (-0.03 logMAR) and increased contrast sensitivity scores (1.69 log CS) on average. This finding coincides with declines in visual function previously measured in healthy ageing populations (Allard et al., 2013; Owsley et al., 1983; Pitts, 1982; Weale, 1975). Though recognition of photo cues was not significantly reduced in older adults (i.e., at ceiling), these visual deficits may have hindered processing of both photos and two-tones. In line with this, poorer contrast sensitivity scores correlated with poorer recognition of photo cues across all participants. However, we found no such correlation for contrast sensitivity and two-tone stimuli, likely due to their high contrast. Instead, poorer visual acuity was found to predict poorer recognition of naive two-tones across both age groups. Together, this suggests that impairment in basic vision accompanying healthy ageing results in less reliable or noisier inputs when viewing both two-tone and photo stimuli. This may not only hinder recognition of these images on initial viewing, but could also affect the integration of information across these two image conditions necessary for perceptual reorganisation. However, no such reductions in perceptual reorganisation were found for older participants. Given the likely disruption to visual input that these basic deficits cause for our older participants, it is striking that we find a preserved cueing effect for these already impoverished stimuli into old age.

Ageing is also associated with declines in some cognitive abilities, which could feasibly affect perceptual reorganisation. While reduced capacity for short-term memory is commonly observed in healthy ageing populations (Cadar et al., 2018; Foster, 2006; Hedden & Gabrieli, 2004; Nilsson, 2003), we did not detect reduced memory of images in our sample of older adults. Here, we specifically tested for memory function that may affect a participant’s ability to perceptually reorganise images in this task. That is, the ability to recall photos of everyday objects over a period of less than 15 minutes. In line with the preservation of working memory of low-level visual features in older populations (Faubert, 2002), we found no reduced performance in older adults’ visual memory for these images compared to younger adults. We can therefore expect that older participants’ ability to retain and recall information from the relevant photo cue was not substantially affected by issues relating to short-term memory.

Ageing may also aid visual perception through accumulated experience and refined long-term knowledge of the visual world. From a Bayesian perspective, when sensory input becomes unreliable, prior knowledge increasingly shapes perceptual inference. In later life, both the decline of low-level visual function - as observed in our older participants - and the lifelong accumulation of visual experience are expected to heighten reliance on such priors (Chan et al., 2021; Moran et al., 2014). Our results support this view. Although older participants showed reduced two-tone recognition, associated with lower visual acuity for naive two-tones, the perceptual benefit of cueing image content was fully preserved and not limited by sensory acuity loss. Indeed, a significant three-way interaction between condition, age, and visual acuity raises the possibility that poorer visual acuity may result in larger cueing benefits in older, but not younger, adults. When sensory input begins to deteriorate with age, the visual system may compensate by placing greater weight on prior knowledge. While replication of this result in larger samples is warranted, this pattern is consistent with a potential protective role of prior knowledge in supporting perceptual performance despite sensory decline as we grow older.

Previous work has suggested that older adults show a stronger influence of prior knowledge when interpreting ambiguous or noisy visual input (Badham et al., 2016; Chan et al., 2021; Loaiza et al., 2015; Wynn et al., 2020). They are less likely to switch percepts when viewing ambiguous stimuli, indicating more stable perceptual interpretations (Botwinick et al., 1959; Dowlati et al., 2016). Neuroimaging evidence links this stability to increased recruitment of frontoparietal and feedback networks, potentially consistent with stronger top-down predictions compensating for weakened sensory precision (Dowlati et al., 2016). Our results fit this pattern: although older adults showed reduced two-tone recognition overall, they exhibited comparable cueing benefits or even enhanced benefits with greater acuity loss, suggesting preserved efficacy, and possibly increased weighting, of prior knowledge during perceptual inference.

This pattern in later life contrasts with early development, where immature feedback mechanisms and limited prior experience may yield weaker perceptual reorganisation (Baum et al., 2020; Milne et al., 2024). Together, these findings support a predictive-coding account in which the precision weighting of sensory input decreases and reliance on priors increases across the lifespan. Similar dynamics have been proposed in clinical populations prone to hallucinations or delusions, where priors are thought to dominate perceptual inference (Davies et al., 2018; Teufel et al., 2015; Zarkali et al., 2019). In healthy ageing, however, knowledge-guided vision appears adaptive rather than pathological, promoting perceptual coherence despite degraded input. In line with this, we recently showed that hallucination proneness was not enhanced in these participants, indicating that unchecked reliance on internally generated percepts is not a feature of healthy ageing (Shenyan et al., 2025).

In summary, we demonstrate that while older adults are less able to recognise impoverished visual scenes compared to younger adults, they receive an equivalent perceptual benefit from informative visual cues. Crucially, this preserved cueing effect occurs despite pronounced age-related declines in basic visual function, which correlated with perception of two-tones before, but not after, exposure to the cue. Within a Bayesian framework, our findings suggest that older adults can compensate for diminished sensory precision by relying on prior knowledge to guide perception. This highlights the resilience of adaptive perceptual mechanisms across the lifespan, including in the presence of sensory decline.

## References

Adámek, P., Langová, V., & Horáček, J. (2022). Early-stage visual perception impairment in schizophrenia, bottom-up and back again. Schizophrenia, 8(1), 27. 10.1038/s41537-022-00237-9

Alderson-Day, B., Smailes, D., Moffatt, J., Mitrenga, K., Moseley, P., & Fernyhough, C. (2019). Intentional inhibition but not source memory is related to hallucination-proneness and intrusive thoughts in a university sample. Cortex; a Journal Devoted to the Study of the Nervous System and Behavior, 113, 267–278. 10.1016/j.cortex.2018.12.020

Allard, R., Renaud, J., Molinatti, S., & Faubert, J. (2013). Contrast sensitivity, healthy aging and noise. Vision Research, 92, 47–52. 10.1016/j.visres.2013.09.004

Andersen, G. J. (2012). Aging and vision: Changes in function and performance from optics to perception. WIREs Cognitive Science, 3(3), 403–410. 10.1002/wcs.1167

Badham, S. P., Hay, M., Foxon, N., Kaur, K., & Maylor, E. A. (2016). When does prior knowledge disproportionately benefit older adults’ memory? Neuropsychology, Development, and Cognition. Section B, Aging, Neuropsychology and Cognition, 23(3), 338–365. 10.1080/13825585.2015.1099607

Bates, D., Mächler, M., Bolker, B., & Walker, S. (2015). Fitting Linear Mixed-Effects Models Using lme4. Journal of Statistical Software, 67(1). 10.18637/jss.v067.i01

Baum, G. L., Cui, Z., Roalf, D. R., Ciric, R., Betzel, R. F., Larsen, B., Cieslak, M., Cook, P. A., Xia, C. H., Moore, T. M., Ruparel, K., Oathes, D. J., Alexander-Bloch, A. F., Shinohara, R. T., Raznahan, A., Gur, R. E., Gur, R. C., Bassett, D. S., & Satterthwaite, T. D. (2020). Development of structure–function coupling in human brain networks during youth. Proceedings of the National Academy of Sciences, 117(1), 771–778. 10.1073/pnas.1912034117

Bona, S., Cattaneo, Z., & Silvanto, J. (2016). Investigating the Causal Role of rOFA in Holistic Detection of Mooney Faces and Objects: An fMRI-guided TMS Study. Brain Stimulation, 9(4), 594–600. 10.1016/j.brs.2016.04.003

Botwinick, J., Robbin, J. S., & Brinley, J. F. (1959). Reorganization Of Perceptions With Age. Journal of Gerontology, 14(1), 85–88. 10.1093/geronj/14.1.85

Cadar, D., Usher, M., & Davelaar, E. J. (2018). Age-Related Deficits in Memory Encoding and Retrieval in Word List Free Recall. Brain Sciences, 8(12), 211. 10.3390/brainsci8120211

Chan, J. S., Wibral, M., Stawowsky, C., Brandl, M., Helbling, S., Naumer, M. J., Kaiser, J., & Wollstadt, P. (2021). Predictive Coding Over the Lifespan: Increased Reliance on Perceptual Priors in Older Adults-A Magnetoencephalography and Dynamic Causal Modeling Study. Frontiers in Aging Neuroscience, 13, 631599. 10.3389/fnagi.2021.631599

Craik, F. I. M. (2023). Memory, aging and the brain: Old findings and current issues. Aging Brain, 4, 100096. 10.1016/j.nbas.2023.100096

Davies, D. J., Teufel, C., & Fletcher, P. C. (2018). Anomalous Perceptions and Beliefs Are Associated With Shifts Toward Different Types of Prior Knowledge in Perceptual Inference. Schizophrenia Bulletin, 44(6), 1245–1253. 10.1093/schbul/sbx177

Davis, S. W., Dennis, N. A., Daselaar, S. M., Fleck, M. S., & Cabeza, R. (2008). Que PASA? The Posterior-Anterior Shift in Aging. Cerebral Cortex, 18(5), 1201–1209. 10.1093/cercor/bhm155

Dowlati, E., Adams, S. E., Stiles, A. B., & Moran, R. J. (2016). Aging into Perceptual Control: A Dynamic Causal Modeling for fMRI Study of Bistable Perception. Frontiers in Human Neuroscience, 10. 10.3389/fnhum.2016.00141

Eppenberger, L. S., Li, C., Wong, D., Tan, B., Garhöfer, G., Hilal, S., Chong, E., Toh, A. Q., Venketasubramanian, N., Chen, C. L.-H., Schmetterer, L., & Chua, J. (2024). Retinal thickness predicts the risk of cognitive decline over five years. Alzheimer’s Research & Therapy, 16(1), 273. 10.1186/s13195-024-01627-0

Faubert, J. (2002). Visual perception and aging. Canadian Journal of Experimental Psychology / Revue Canadienne de Psychologie Expérimentale, 56(3), 164–176. 10.1037/h0087394

Flounders, M. W., González-García, C., Hardstone, R., & He, B. J. (2019). Neural dynamics of visual ambiguity resolution by perceptual prior. eLife, 8, e41861. 10.7554/eLife.41861

Foster, T. C. (2006). Biological markers of age-related memory deficits: Treatment of senescent physiology. CNS Drugs, 20(2), 153–166. 10.2165/00023210-200620020-00006

Friston, K., & Kiebel, S. (2009). Predictive coding under the free-energy principle. Philosophical Transactions of the Royal Society of London. Series B, Biological Sciences, 364(1521), 1211–1221. 10.1098/rstb.2008.0300

González-García, C., Flounders, M. W., Chang, R., Baria, A. T., & He, B. J. (2018). Content-specific activity in frontoparietal and default-mode networks during prior-guided visual perception. eLife, 7, e36068. 10.7554/eLife.36068

Hardstone, R., Zhu, M., Flinker, A., Melloni, L., Devore, S., Friedman, D., Dugan, P., Doyle, W. K., Devinsky, O., & He, B. J. (2021). Long-term priors influence visual perception through recruitment of long-range feedback. Nature Communications, 12, 6288. 10.1038/s41467-021-26544-w

Hedden, T., & Gabrieli, J. D. E. (2004). Insights into the ageing mind: A view from cognitive neuroscience. Nature Reviews Neuroscience, 5(2), 87–96. 10.1038/nrn1323

Hsieh, P.-J., Vul, E., & Kanwisher, N. (2010). Recognition Alters the Spatial Pattern of fMRI Activation in Early Retinotopic Cortex. Journal of Neurophysiology, 103(3), 1501–1507. 10.1152/jn.00812.2009

Imamoglu, F., Kahnt, T., Koch, C., & Haynes, J.-D. (2012). Changes in functional connectivity support conscious object recognition. NeuroImage, 63(4), 1909–1917. 10.1016/j.neuroimage.2012.07.056

Kéri, S., Antal, A., Szekeres, G., Benedek, G., & Janka, Z. (2002). Spatiotemporal Visual Processing in Schizophrenia. The Journal of Neuropsychiatry and Clinical Neurosciences, 14(2), 190–196. 10.1176/jnp.14.2.190

Lai, L. Y., Frömer, R., Festa, E. K., & Heindel, W. C. (2020). Age-related changes in the neural dynamics of bottom-up and top-down processing during visual object recognition: An electrophysiological investigation. Neurobiology of Aging, 94, 38–49. 10.1016/j.neurobiolaging.2020.05.010

Lee, T. S., & Mumford, D. (2003). Hierarchical Bayesian inference in the visual cortex. Journal of the Optical Society of America A, 20(7), 1434. 10.1364/JOSAA.20.001434

Loaiza, V. M., Rhodes, M. G., & Anglin, J. (2015). The Influence of Age-Related Differences in Prior Knowledge and Attentional Refreshing Opportunities on Episodic Memory. The Journals of Gerontology Series B: Psychological Sciences and Social Sciences, 70(5), Article 5. 10.1093/geronb/gbt119

Martin, D., Fowlkes, C., Tal, D., & Malik, J. (2001). A database of human segmented natural images and its application to evaluating segmentation algorithms and measuring ecological statistics. Proceedings Eighth IEEE International Conference on Computer Vision. ICCV 2001, *2*, 416–423. 10.1109/ICCV.2001.937655

Mazuz, Y., Kessler, Y., & Ganel, T. (2024). Age-related changes in the susceptibility to visual illusions of size. Scientific Reports, 14(1), 14583. 10.1038/s41598-024-65405-6

Milne, G. A., Lisi, M., McLean, A., Zheng, R., Groen, I. I. A., & Dekker, T. M. (2024). Perceptual reorganization from prior knowledge emerges late in childhood. iScience, 27(2), 108787. 10.1016/j.isci.2024.108787

Mooney, C. M. (1957). Age in the development of closure ability in children. Canadian Journal of Psychology, 11(4), 219–226. 10.1037/h0083717

Moran, R. J., Symmonds, M., Dolan, R. J., & Friston, K. J. (2014). The Brain Ages Optimally to Model Its Environment: Evidence from Sensory Learning over the Adult Lifespan. PLOS Computational Biology, 10(1), e1003422. 10.1371/journal.pcbi.1003422

Nagarajan, N., Assi, L., Varadaraj, V., Motaghi, M., Sun, Y., Couser, E., Ehrlich, J. R., Whitson, H., & Swenor, B. K. (2022). Vision impairment and cognitive decline among older adults: A systematic review. BMJ Open, 12(1), e047929. 10.1136/bmjopen-2020-047929

Nilsson, L.-G. (2003). Memory function in normal aging. Acta Neurologica Scandinavica, 107(s179), 7–13. 10.1034/j.1600-0404.107.s179.5.x

Owsley, C. (2011). Aging and vision. Vision Research, 51(13), 1610–1622. 10.1016/j.visres.2010.10.020

Owsley, C., Sekuler, R., & Siemsen, D. (1983). Contrast sensitivity throughout adulthood. Vision Research, 23(7), 689–699. 10.1016/0042-6989(83)90210-9

Pitts, D. G. (1982). Visual acuity as a function of age. Journal of the American Optometric Association, 53(2), 117–124.

Rauss, K., & Pourtois, G. (2013). What is Bottom-Up and What is Top-Down in Predictive Coding? Frontiers in Psychology, 4. https://www.frontiersin.org/article/10.3389/fpsyg.2013.00276

Reyes-Ortiz, C. A., Kuo, Y., DiNuzzo, A. R., Ray, L. A., Raji, M. A., & Markides, K. S. (2005). Near Vision Impairment Predicts Cognitive Decline: Data from the Hispanic Established Populations for Epidemiologic Studies of the Elderly. Journal of the American Geriatrics Society, 53(4), 681–686. 10.1111/j.1532-5415.2005.53219.x

Sheldon, S., Pichora-Fuller, M. K., & Schneider, B. A. (2008). Priming and sentence context support listening to noise-vocoded speech by younger and older adults. The Journal of the Acoustical Society of America, 123(1), 489–499. 10.1121/1.2783762

Shenyan, O., Haye, L., Milne, G. A., Lisi, M., Greenwood, J. A., Skipper, J. I., & Dekker, T. M. (2025). Reduced susceptibility to experimentally-induced complex visual hallucinations with age. Cortex, 191, 188–204. 10.1016/j.cortex.2025.08.001

Shoham, N., Lewis, G., Hayes, J. F., Silverstein, S. M., & Cooper, C. (2023). Association between visual impairment and psychosis: A longitudinal study and nested case-control study of adults. Schizophrenia Research, 254, 81–89. 10.1016/j.schres.2023.02.017

Snodgrass, J. G., & Vanderwart, M. (1980). A standardized set of 260 pictures: Norms for name agreement, image agreement, familiarity, and visual complexity. Journal of Experimental Psychology: Human Learning and Memory, 6(2), 174–215. 10.1037/0278-7393.6.2.174

Teufel, C., Subramaniam, N., Dobler, V., Perez, J., Finnemann, J., Mehta, P. R., Goodyer, I. M., & Fletcher, P. C. (2015). Shift toward prior knowledge confers a perceptual advantage in early psychosis and psychosis-prone healthy individuals. Proceedings of the National Academy of Sciences of the United States of America, 112(43), 13401–13406. 10.1073/pnas.1503916112

The MathWorks Inc. (2015). MATLAB version: 8.6.0.267246 (R2015b) [Computer software]. The MathWorks Inc. https://www.mathworks.com

The MathWorks Inc. (2022). MATLAB version: 9.12.0.1956245 (R2022a) [Computer software].

The MathWorks Inc. https://www.mathworks.com

van Loon, A. M., Fahrenfort, J. J., van der Velde, B., Lirk, P. B., Vulink, N. C. C., Hollmann, M. W., Scholte, H. S., & Lamme, V. A. F. (2016). NMDA Receptor Antagonist Ketamine Distorts Object Recognition by Reducing Feedback to Early Visual Cortex. Cerebral Cortex (New York, N.Y.: 1991), 26(5), 1986–1996. 10.1093/cercor/bhv018

Weale, R. A. (1975). Senile changes in visual acuity. Transactions of the Ophthalmological Societies of the United Kingdom, 95(1), 36–38.

Whitson, H. E., Cronin-Golomb, A., Cruickshanks, K. J., Gilmore, G. C., Owsley, C., Peelle, J. E., Recanzone, G., Sharma, A., Swenor, B., Yaffe, K., & Lin, F. R. (2018). American Geriatrics Society and National Institute on Aging Bench-to-Bedside Conference: Sensory Impairment and Cognitive Decline in Older Adults. Journal of the American Geriatrics Society, 66(11), 2052–2058. 10.1111/jgs.15506

Wynn, J. S., Ryan, J. D., & Moscovitch, M. (2020). Effects of prior knowledge on active vision and memory in younger and older adults. Journal of Experimental Psychology: General, 149(3), 518–529. 10.1037/xge0000657

Zarkali, A., Adams, R. A., Psarras, S., Leyland, L.-A., Rees, G., & Weil, R. S. (2019). Increased weighting on prior knowledge in Lewy body-associated visual hallucinations. Brain Communications, 1(1), fcz007. 10.1093/braincomms/fcz007

Zheng, D. D., Swenor, B. K., Christ, S. L., West, S. K., Lam, B. L., & Lee, D. J. (2018). Longitudinal Associations Between Visual Impairment and Cognitive Functioning: The Salisbury Eye Evaluation Study. JAMA Ophthalmology, 136(9), 989. 10.1001/jamaophthalmol.2018.2493

